# AIP56, an AB toxin secreted by *Photobacterium damselae* subsp. *piscicida*, has tropism for myeloid cells

**DOI:** 10.1101/2024.11.12.623249

**Authors:** Inês Lua Freitas, Fátima Macedo, Liliana Oliveira, Pedro Oliveria, Ana do Vale, Nuno M.S. dos Santos

## Abstract

The AB-type toxin AIP56 is a key virulence factor of *Photobacterium damselae* subsp. *piscicida* (*Phdp*), inducing apoptosis in fish immune cells. The discovery of AIP56-like and AIP56-related toxins in diverse organisms, including human-associated *Vibrio* strains, highlights the evolutionary conservation of this toxin family, suggesting that AIP56 and its homologs may share conserved receptors across species. These toxins have potential for biotechnological applications, such as therapeutic protein delivery and immune modulation. Herein, the cell specificity of AIP56 for immune cells in sea bass, mice, and humans was characterized. Whether AIP56 interacts directly or indirectly with sea bass neutrophils was never investigated and it was shown that only a small population of sea bass neutrophils internalized AIP56, indicating that most of the neutrophilic destruction during *Phdp* infection and/or AIP56 intoxication does not result from the direct toxicity of the toxin. Moreover, the cellular tropism of AIP56 for myeloid cells was observed in the three species, including its preference for macrophages. Further, mouse and human M0 and M2-like macrophages internalized more toxin than M1-like macrophages. Despite the limited interaction of lymphoid cells with AIP56, mouse B1-cells were able to internalize the toxin, possibly due to its myeloid features. These findings are relevant for both pathogenicity and biomedical contexts.

## 1 Introduction

*Photobacterium damselae* subsp. *piscicida* (*Phdp*) is a Gram-negative bacterium that infects a large number of warm saltwater fish species, including sea bass, sea bream, sole and yellowtail, with high impact for the aquaculture industry (1–3). A crucial virulence factor of this pathogen is AIP56 (Apoptosis-Inducing Protein of 56 kDa), a plasmid-encoded AB-type toxin secreted through T2SS (type 2 secretion system) (4–6). It has been reported that the systemic distribution of the toxin in infected animals is associated to the death of fish macrophages and neutrophils by post-apoptotic secondary necrosis, leading to the release of highly cytotoxic molecules that likely contribute to the typical necrotic lesions of *Phdp* infections (4,7–9). The AIP56-associated systemic elimination of the host phagocytes facilitates the extracellular proliferation of the pathogen, leading to septicemia and death of the infected host (8). Passive immunization with rabbit anti-AIP56 serum protected sea bass from *Phdp* infections, further supporting that AIP56 is a key virulence factor of this pathogen (4). Although *Phdp* is unable to grow at 37°C and has a strict salt dependency (4), making it incapable to infect mammals, AIP56 displays high toxicity for mouse macrophages (10).

The three-dimensional structure of AIP56 has been recently solved, showing that it has a three-domain organization (11). The N-terminal domain is homologous to NleC (non LEE-encoded effector C), a T3SS effector secreted by several enteric bacteria associated with human diseases (12–14) and displays metalloprotease activity for NF-κB p65 (6), a transcription factor that promotes the expression of anti-apoptotic and inflammatory genes (15). Thus, AIP56-mediated cleavage of NF-κB p65 will likely lead to decreased expression of anti-apoptotic genes and consequently apoptotic death of the host cells (6,8,10). The small middle domain is mostly composed by a 35-aa long partially structured linker peptide, whose ends are bound by a disulfide bridge necessary for intoxication (10), and is likely involved in transmembrane channel formation (11). The C-terminal domain is responsible for receptor-binding (6) and is also required for pore-formation (11). Studies performed in sea bass and mouse macrophages showed that to reach its molecular target in the cytosol, AIP56 binds to an unknown cell surface receptor(s) and is internalized by clathrin-dependent endocytosis followed by acidic-dependent translocation from the endosomal compartment into the cytosol, in a process involving Hsp90 and Cyclophilin A/D (6,10,16).

An increasing number of putative toxins homologous to AIP56 (AIP56-like toxins) is being annotated in the genomes of organisms that have hosts from different phylogenetic groups such as *Shewanella psychrophila, Arsenophonus nasoniae, Candidatus* Symbiopectobacterium and several *Vibrio* species, including strains that have been isolated from human blood and stool (17,18). Also, toxins that have a domain homologous to the receptor-binding domain of AIP56 but distinct catalytic and middle domains (AIP56-related toxins) can be found in the genomes of prokaryotic and eukaryotic organisms such as *Enterovibrio norvegicus, Shewanella woodyi, Shewanella nanhaiensis, Neisseria gonorrhoeae, Danaus plexippus, Danaus chrysippus, Operophtera brumata, Thrips palmi, Drosophila ananassae* (19) and *Drosophila bipectinata*. Moreover, proteins homologous to the AIP56’s receptor-binding domain are also present in the genome of *Acyrthosiphon pisum* secondary endosymbiont 2 (APSE2) and 7 (APSE7) bacteriophages. AlphaFold predictions indicate that the homologous domain present in all these putative toxins are structurally similar to the receptor-binding domain of AIP56 (11), which, together with the fact that AIP56 is toxic to both sea bass and mouse macrophages (10,16), suggests that they likely bind to a phylogenetically conserved receptor and target similar cell populations.

In this work, a detailed characterization of the specificity of AIP56 for cells involved in the immune response in sea bass, humans and mice was performed. Identification of AIP56 target cells is not only important for understanding the biology of intoxication, with eventual extension to other members of the AIP56-toxin family, but also relevant from the biotechnological and/or biomedical point of view. In fact, AB toxins naturally evolved as autonomous systems capable of delivering their catalytic domain into the cytosol of target cells and are top candidates for delivering therapeutic proteins to the cytosol of human cells to treat many diseases and conditions (19,20). The modular structure of these toxins allows replacing their domains or grafting other moieties to diversify their functions or redirect them to different cell types. In the case of AIP56, it has been shown that the delivery region (middle and receptor-binding domains) is capable of delivering β-lactamase into the cytosol of mouse macrophages (11,16). The here reported characterization of the cellular susceptibility to AIP56 in humans (and mice, as a model species) is of relevance when considering the future use of AIP56 and AIP56-like and/or -related toxins as biotechnological/biomedical tools.

## 2 Results

### 2.1 AIP56 internalization by sea bass leukocytes

The occurrence of apoptosis of macrophages and neutrophils after injection of sea bass with *Phdp* or AIP56 is well documented (4,8), but the causal correlation with AIP56 was only demonstrated for macrophages (10,21). Thus, it was important to evaluate if AIP56 is also able to directly target other sea bass immune cells, namely neutrophils and lymphocytes.

Sea bass neutrophils can hardly be manipulated outside the host as they die quickly. To circumvent this, the internalization of AIP56 in sea bass neutrophils was studied in vivo, in inflamed peritoneal cavities. UV-killed *Phdp* MT1415 cells were injected intraperitoneally (ip) to attract neutrophils to the peritoneal cavity and, after 6 h, fluorescent-labelled AIP56 (AIP56-488) was ip injected in the fish. After 15 min, peritoneal cells were collected and the percentage of AIP56-488-positive neutrophils and macrophages analyzed by flow cytometry. The populations of gated neutrophils and macrophages analyzed were selected based on their high side (cell complexity) and forward (cell size) scatter profiles, typical of neutrophils and macrophages. Cell sorting and positive or negative detection of intracellular peroxidase activity (22) confirmed that > 95% of the gated cells were neutrophils and > 99% were macrophages, respectively (Figure S1). Approximately 82% of macrophages and 13 % of neutrophils were positive for AIP56-488 (Figure 1; Supplementary Table 1), showing that AIP56 is internalized by the majority of macrophages, but only by a minoritarian population of neutrophils. A population of small mononucleated cells was also observed in these sea bass inflamed peritoneal cavities, which is composed by lymphocytes and thrombocytes (23). No AIP56-488 positive cells were identified in this population (data not shown). Lymphocytes and thrombocytes cannot be distinguished by flow cytometry based on their cell complexity and size (23), therefore, to study the internalization of AIP56 by sea bass lymphocytes, cell suspensions enriched in lymphocytes were obtained from sea bass spleen or thymus by centrifugation on a Percoll gradient. Cell suspensions were then incubated with AIP56-488 for 15 min on ice followed by 15 min at 22 °C to allow toxin internalization (10,21), and the percentage of AIP56-488 positive cells analyzed by flow cytometry using anti-sea bass IgM antibody (21) to identify splenic IgM^+^ B-cells and thymic IgM^-^ cells (putatively thymocytes, since ∼ 74.9% thymus IgM^-^ cells are thymocytes (24–26)). After incubation with AIP56, only a very small percentage of AIP56-488 positive IgM^+^ B-cells was detected and the MFI of AIP56-488-positive cells was low, indicating that most sea bass lymphoid cells appear unable to internalize the toxin.

**Figure 1.**
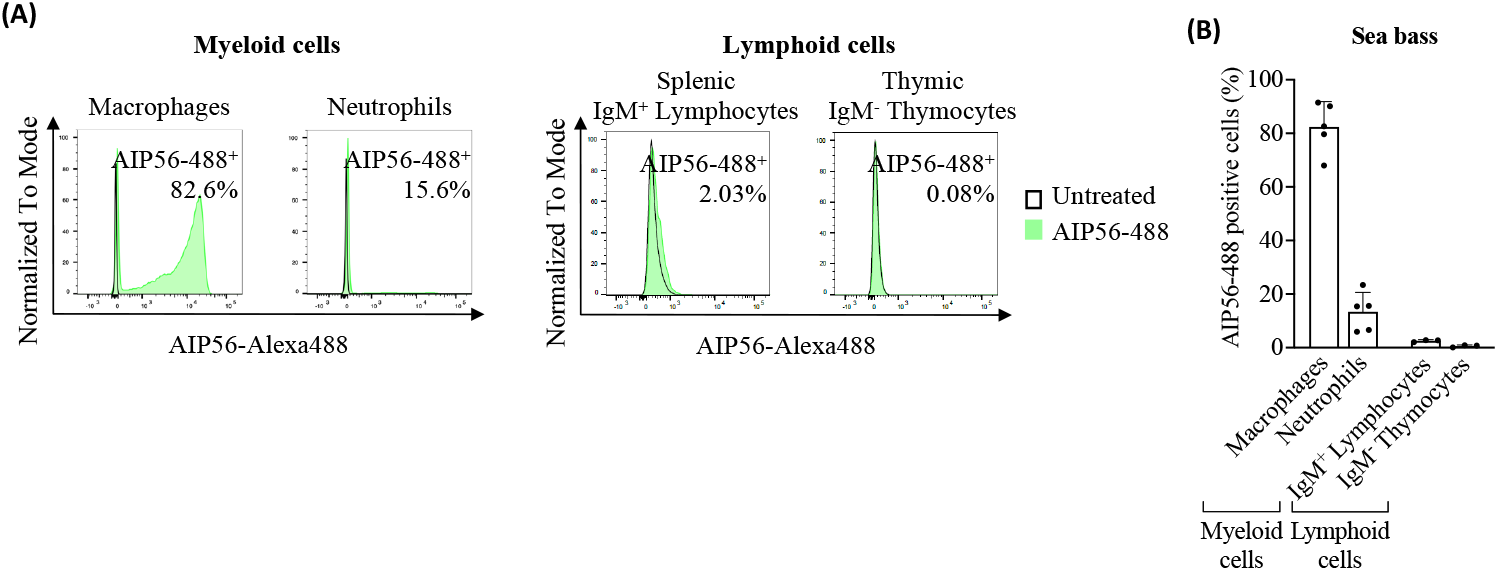
Internalization of AIP56 by sea bass cells. **(A)** Representative flow cytometry plots and **(B)** quantification of AIP56-488 internalization by sea bass macrophages (n=5), neutrophils (n=5) and lymphoid cells (n=3). Macrophages and neutrophils were collected from sea bass inflamed peritoneal cavity 15 min after injection of AIP56-488. Lymphoid cells obtained from sea bass spleen or thymus were left untreated or incubated with AIP56-488 for 15 min on ice followed by 15 min at 22 °C. The percentage of AIP56-488-positive cells was quantified by flow cytometry after gating the different cell populations (Figure S1). Flow cytometry source data is shown in Supplementary Material data sheet 1.

### 2.2 AIP56 internalization by mouse leukocytes

Given the existence of a putative AIP56-like toxin in *Vibrio tarriae* isolated from humans (17,18) and the AIP56 and AIP56-like and -related toxins potential as biotechnological/biomedical tools, the study was extended to mouse and human immune cells.

Mouse myeloid cells were obtained from bone marrow (BM) (macrophages, monocytes, eosinophils and neutrophils), spleen (macrophages, monocytes and dendritic cells (DCs)), peritoneum (macrophages and neutrophils) and blood (neutrophils). Spleen was also the source of lymphoid cells (B-cells and T-cells). The cells were incubated for 15 min on ice with AIP56-488, followed by 30 min at 37 °C to allow toxin internalization (10,27). Mouse cell types were identified with established fluorescent cell markers (Figure S2-S4) and the percentage of AIP56-488-labeled cells analyzed by flow cytometry.

The results (Figure 2A and B; Supplementary Table 2) showed that on average 73 to 98 % of macrophages became positive for AIP56-488, regardless of whether they were obtained from the BM, spleen or peritoneum. More than 20% of BM and spleen monocytes as well as DCs from spleen also became positive for AIP56-488. Moreover, the MFI in AIP56-488 positive monocytes and DCs was high, and even higher in macrophages (Supplementary Table 2). Less than 4% of mouse BM eosinophils and neutrophils, around 9% and 7% of peritoneal and blood neutrophils, respectively, were positive for AIP56-488, although the MFI of these cells was lower than that in AIP56-488 positive monocytes, DCs and macrophages (Figure 2B and Supplementary Table 2).

**Figure 2.**
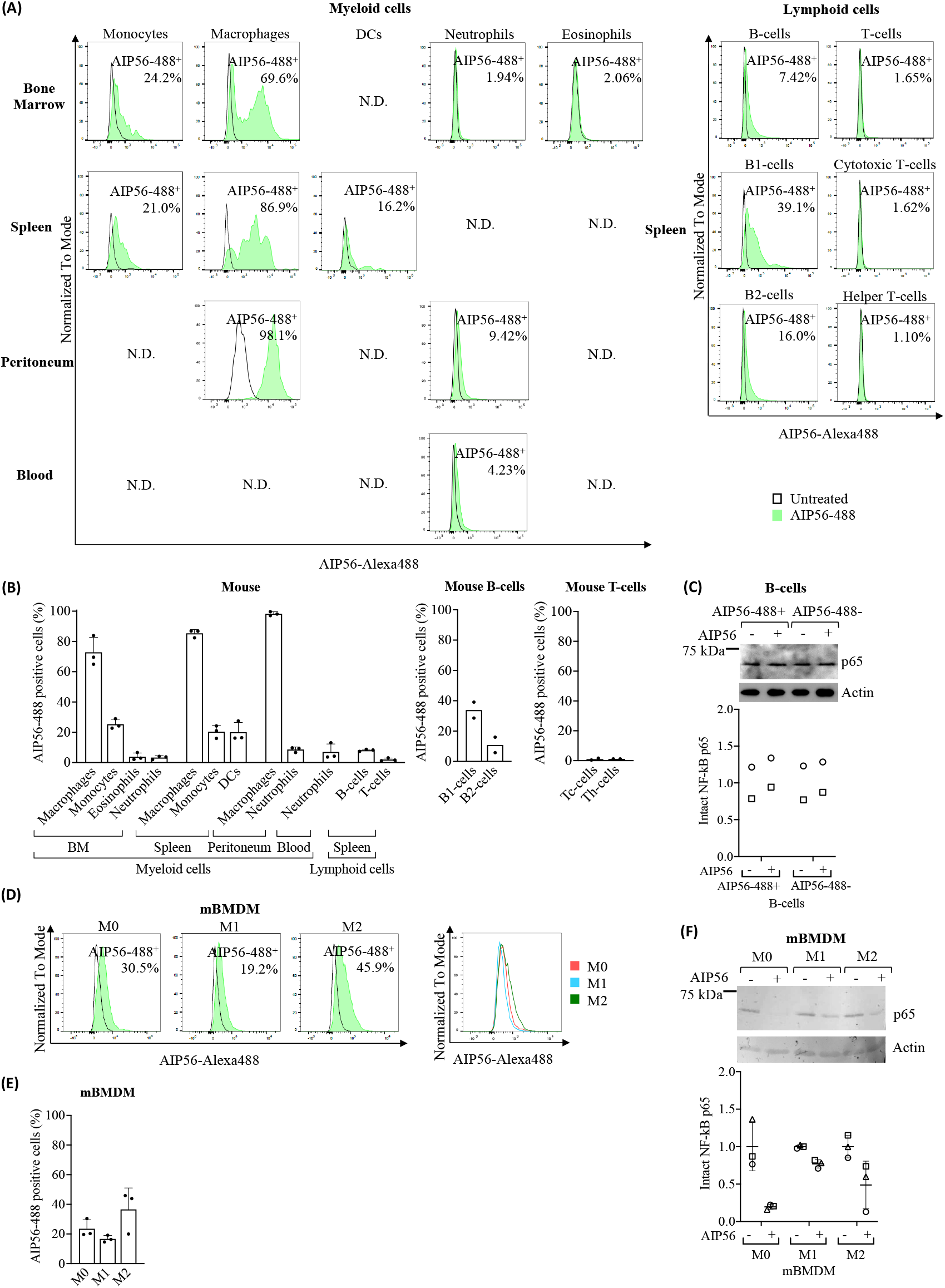
Internalization of AIP56 by mouse leukocytes. **(A)** Representative flow cytometry plots (N.D.: not done) and **(B)** quantification of AIP56-488 internalization by mouse myeloid (n=3) and lymphoid cells (n=3, except B1-cells, B2-cells, cytotoxic T-cells (Tc-cells) and helper T-cells (Th-cells): n=2). B1-cells, B2-cells, Tc-cells and helper Th-cells were analyzed from spleen. **(C)** Representative WB analysis of NF-κB p65 cleavage in mouse AIP56-488+ and AIP56-488? B-cells intoxicated with AIP56 (n=2). **(D)** Representative flow cytometry plots of AIP56 internalization by mBMDM polarized into M0, M1-like or M2-like phenotypes and comparison of representative flow cytometry plots of AIP56-488 positive cells (overlay plot, right panel). **(E)** Quantification of AIP56 internalization by mBMDM polarized into M0, M1-like or M2-like phenotypes (n=3) **(F)** Representative WB analysis of NF-κB p65 cleavage in M0, M1-like or M2-like mBMDM incubated with AIP56 (n=3). For toxin internalization **(A, B, D** and **E)** cells were left untreated or incubated with AIP56-488 for 15 min on ice followed by 30 min at 37 °C and the percentage of AIP56-488 positive cells quantified by flow cytometry after gating the different cell populations with specific cell markers (Figure<u>s</u> S2-S4). For NF-κB p65 cleavage **(C, F)**, cells were incubated with AIP56 for 15 min on ice followed by 4 h at 37 °C and quantification of intact NF-κB p65 bands; loading correction was achieved by dividing the density of p65 band by the density of actin band from the same sample; results are presented as fold-change relative to the untreated control. Different symbols represent independent experiments. Flow cytometry and WB source data is shown in Supplementary Material data sheet 1.

Regarding mouse lymphocytes (Figure 2A and B; Supplementary Table 2), approximately 8% of AIP56-488 positive B-cells were detected, being the percentage of positive B1-cells (∼ 34%) three times higher than the percentage of positive B2-cells (∼ 11%). In turn, almost no internalization of AIP56-488 was detected in mouse T-cells, with less than 1.11% cytotoxic T-cells and helper T-cells becoming positive for the toxin. These results suggest that there is a small population of B cells that internalize AIP56. To investigate whether the internalization of AIP56 by B-cells (∼ 8%) leads to NF-κB p65 cleavage, mouse B-cells were incubated with inactive AIP56-488 and the B-cells positive for inactive AIP56-488 were sorted and subsequently incubated with wild-type AIP56 to evaluated p65 cleavage by Western Blotting (WB). However, p65 cleavage was not detected when AIP56-488 positive B-cells were incubated with AIP56 (Figure 2C). It remains to be investigated if this results from a defective translocation or refolding mechanism of AIP56 in the cytosol of these cells or to other, so far unknown, specificities of these cells.

Considering the preference of AIP56 to intoxicate macrophages, and the distinct role of macrophage subpopulations in the immune response (28–31) the susceptibility of M0 (non-activated/resting), M1-like (classically activated/proinflammatory) and M2-like (alternatively activated/anti-inflammatory) mBMDM to AIP56 was investigated by assessing both internalization (Figure 2D and E; Supplementary Table 2) and p65 cleavage (Figure 2F). Interestingly, there appears to be a preference for the internalization of AIP56 in M0 macrophages and, especially, in M2-like, when compared to M1-like macrophages (Figure 2D and E; Supplementary Table 2). Accordingly, the cleavage of p65 was less pronounced in M1-like macrophages (Figure 2F), especially compared to M0-like macrophages.

### 2.3 AIP56 internalization by human leukocytes

Human monocytes and B and T lymphocytes were isolated from human buffy coats. Monocytes were directly used or differentiated to M0, M1-like or M2-like macrophages, or monocyte-derived dendritic cells (moDCs). Human neutrophils were obtained from peripheral blood. The percentages of AIP56-488-labeled cells were analyzed essentially as described for mouse cells, but using human cell markers for cell type identification (Figures S5 and S6).

The results of the internalization assay covering myeloid cells showed that on average over 92% of monocytes, M0, M1-like and M2-like macrophages, became positive for AIP56-488, and more than 72% of moDCs internalized the toxin (Figure 3A and B; Supplementary Table 3). Although the percentages of AIP56-488 positive cells did not differ among the different macrophage phenotypes, the MFI was higher in M0 and M2 macrophages, suggesting that these cell subtypes internalize the toxin with higher efficiency. However, no differences in the AIP56-induced cleavage of p65 among human macrophages subtypes were detected. Cleavage of NF-κB p65 was also detected in monocytes and moDCs incubated with AIP56 (Figure 3C). Finally, contrary to what was observed in mice, a large percentage of human neutrophils (∼ 61%) internalized AIP56-488 (Figure 3A and B; Supplementary Table 3).

**Figure 3.**
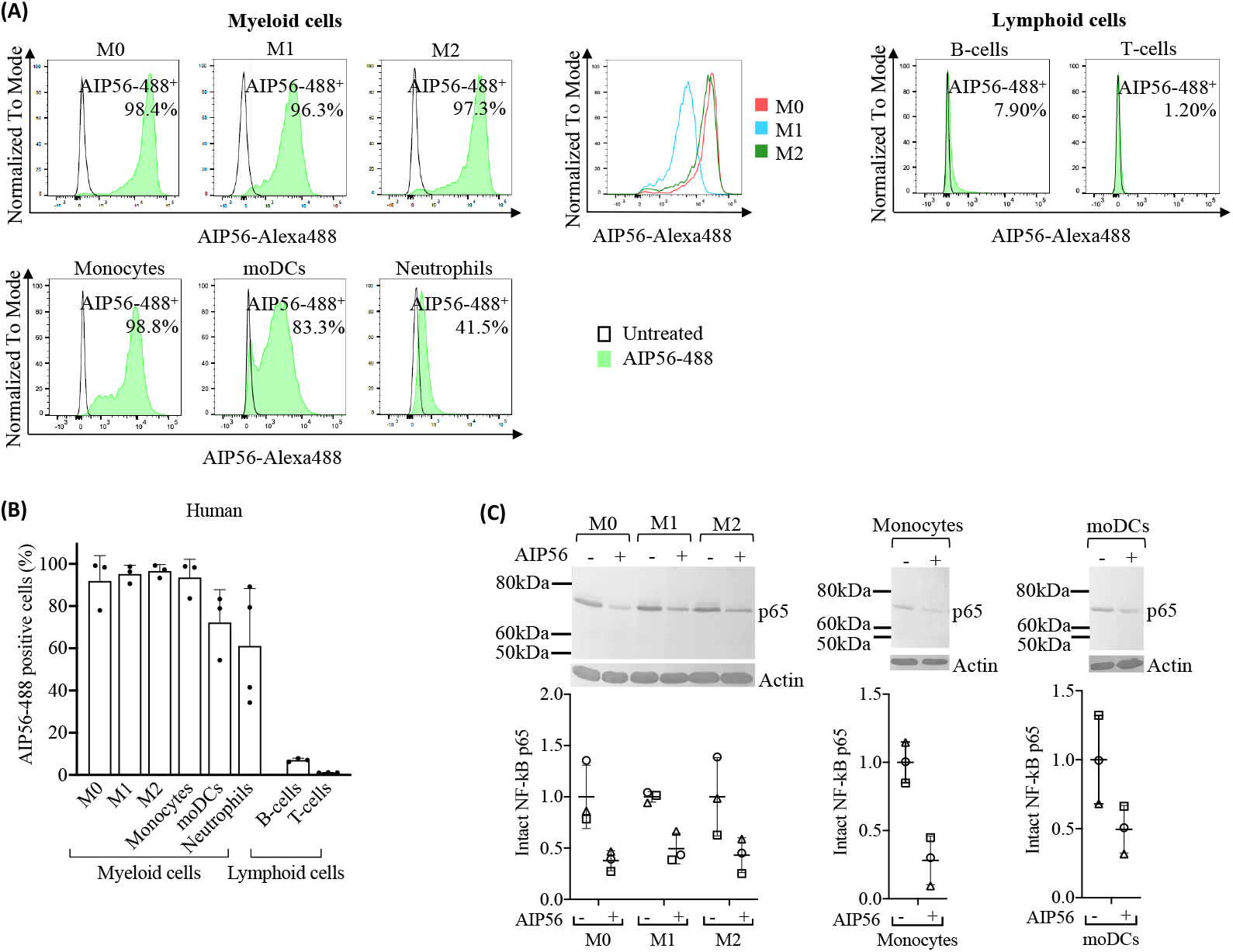
Internalization of AIP56 by human leukocytes. **(A)** Representative flow cytometry plots of AIP56-488 internalization by human myeloid and lymphoid cells and comparison of representative flow cytometry plots of AIP56-488 positive cells from human monocyte derived macrophages M0 (red), M1-like (blue), M2-like (green). **(B)** Quantification of AIP56-488 internalization by human myeloid (n=3 except neutrophils, for which n=4) and lymphoid cells (n=3). For assessing internalization, leukocytes were left untreated or incubated with AIP56-488 for 15 min on ice followed by 30 min at 37 °C and the percentage of AIP56-488-positive cells quantified by flow cytometry after gating the different cell populations with specific cell markers (Figures S5 and S6). **(C)** Representative WB analysis of NF-κB p65 cleavage in human M0, M1-like or M2-like macrophages, monocytes and moDCs incubated with AIP56 for 15 min on ice followed by 6 h (macrophages) or 4 h (monocytes and moDCs) at 37 °C and quantification of intact NF-κB p65 bands (n=3). Loading correction was achieved by dividing the density of p65 band by the density of actin band from the same sample. Results are presented as fold-change relative to the untreated control. Different symbols represent independent experiments. Flow cytometry and WB source data is shown in Supplementary Material data sheet 1.

Regarding human lymphocytes (Figure 3A and B; Supplementary Table 3), a similar result to that obtained for mouse lymphocytes was observed, with ∼ 7% of B-cells positive for AIP56-488 and almost no AIP56-488 positive T-cells (∼ 1%). Nevertheless, the MFI of AIP56-488 positive B- and T-cells was very low, indicating low internalization of the toxin in these cells.

## 3 Discussion

Sea bass is one of the main fish species targeted by *Phdp* virulent strains and has been widely used experimentally to study the virulence factors and mechanisms behind the disease caused by this pathogen. Pathogenic strains of *Phdp* secrete AIP56 in large quantities, which plays a crucial role in the virulence and pathological hallmarks observed during the disease (4–6,8). Among these, the elimination of macrophages and neutrophils by post-apoptotic secondary necrosis stands out, disarming the host’s innate immune response against the pathogen and contributing to the occurrence of extensive tissue destruction (4,8,9). However, while the direct intoxication of macrophages by AIP56 has been well documented (6,10,21), its direct effect on neutrophils and other immune cells remained to be investigated.

In this work, it was shown that sea bass macrophages internalize AIP56 with high efficiency, corroborating their elimination by the toxin in infected animals (4,8,9). In contrast, despite the extensive neutrophil apoptosis observed during *Phdp* infection (8,9) or AIP56 intoxication (4,8), only a small subset of neutrophils became positive for AIP56 after incubation with the toxin, indicating that most of the observed neutrophil destruction during *Phdp* infection may not result from the direct toxicity of AIP56. Instead, it may be a consequence of cell death pathways that occur during the participation of neutrophils in the inflammatory response (32,33) triggered by *Phdp* or its components. In the absence of live macrophages to clear dying neutrophils, they lyse and release highly cytotoxic molecules that likely contribute to the extensive necrotic lesions typically found in *Phdp* infected fish (4,7–9).

The availability of monoclonal antibodies against sea bass IgM (34) also allowed to evaluate the internalization of AIP56 in lymphoid cells. The fact that only a very low percentage of B-cells became positive for AIP56, with relatively low MFI, and almost no AIP56-4886 positive IgM^-^ thymocytes were observed, suggests that sea bass lymphocytes lack or have low levels of the receptor(s) for the toxin, supporting why no significant alterations in these cell types have been reported during *Phdp* infection or AIP56 intoxication.

Altogether, these data show that, in sea bass, AIP56 targets myeloid cells, having greater tropism for macrophages. The myeloid lineage is an essential component of the innate immune response, playing a key role in the initial response against infections (35,36). Furthermore, myeloid cells bridge innate and adaptive immunity through antigen processing and presentation, mainly by dendritic cells and macrophages, thus triggering the adaptive immune response (35,36). Thus, by disabling key cells of the myeloid lineage, AIP56 evades both arms of the immune response, facilitating bacterial proliferation and subsequent septicemia and death of the host (8,37).

As observed in sea bass, the results also indicate that AIP56 has specificity for human and mouse myeloid cells, preferably macrophages. In mice, there appears to be a greater tropism for peritoneal macrophages (AIP56-positive cells), accompanied by greater toxin content (MFI), compared to those in the spleen and BM. Peritoneal macrophages have been described as the most mature, followed by splenic and then BM macrophages (38). In the present study, peritoneal macrophages were obtained 6 h after proinflammatory stimulation. In this condition, it has been described that resident peritoneal macrophages are in an active/mature sate and a small population of peritoneal macrophages derive from recruited circulating monocytes that, in later stages of the inflammation, eliminate resident macrophages and acquire mature resident identity (39–41). Thus, the preference of AIP56 may depend on the maturation/activation status of macrophages, which likely correlates with differential expression of AIP56 receptor(s). Furthermore, it may also correlate with the monocytic origin of some of the peritoneal macrophages present after the proinflammatory stimuli. Indeed, in humans, it was observed that AIP56 has high affinity not only for macrophages but also for dendritic cells, which have been both derived from circulating monocytes.

Regarding the affinity of AIP56 for classically categorized macrophage subpopulations (42), there appears to be a preference to M0- (resting) and M2-like (anti-inflammatory) over M1-like (pro-inflammatory) macrophages. This is mostly supported by results in mice, where a lower percentage of AIP56-positive cells and weaker p65 cleavage was observed in M1-like macrophages. On the other hand, in humans only a greater internalization efficiency (higher MFI) was observed in these macrophage subsets. Considering that the macrophage subsets were derived ex vivo from different cellular compartments, it is not possible to discern whether the observed differences are species specific or actually result from their distinct origin. Although the existence of subpopulations equivalent to mammalian M1- and M2-like macrophages has been described in fish (43), the lack of specific antibodies to distinguish these subpopulations did not allow to investigate their susceptibility to AIP56 and the consequences for infection.

In mice, in agreement to what was observed in sea bass, only a low percentage of neutrophils (<10 %) internalized AIP56 regardless of their source (BM, blood and peritoneum). As in blood neutrophils, higher internalization appears to occur in peritoneal neutrophils recruited after inflammatory stimulation, which suggests greater expression of the AIP56 receptor(s) in activated neutrophils (44). In contrast to what was observed in mice, a much higher percentage of human blood neutrophils (61 %) internalized AIP56. This, indicates species-specific differences in the expression of the AIP56 receptor(s), which agrees with the documented differences between human and mice neutrophils, including a distinct receptor repertoire (33).

Finally, some mouse and human B lymphocytes (7-8%) internalized AIP56. Interestingly, the results obtained in mice show that the toxin has greater affinity for B1 lymphocytes, which express transcription factors and surface markers that confers them myeloid characteristics (45,46). In fact, while B2 lymphocytes originate from the bone marrow, mature in the blood and secondary lymphatic organs, and play an essential role in the adaptive immune response by producing memory cells and high-affinity antibodies (47,48), B1 lymphocytes originate from fetal liver progenitors, have self-renewal capacity, and play a role in the innate immune response, including phagocytic and antigen presentation capabilities (45,47,49). However, AIP56 internalization in B-cells did not result in detectable NF-κB p65 cleavage, suggesting that AIP56 may not be fully functional in these cells or that additional factors are required for its activity.

Overall, it can be concluded that the efficient internalization of AIP56 by sea bass, mouse and human macrophages and subsequent cleavage of NF-κB p65 points to the evolutionary conservation of its receptor(s) and mechanism of action across species. This finding is relevant as it raises the possibility that AIP56-like and -related toxins may also play a role in pathogenesis, with particular interest in the context of the AIP56-like toxin identified in human-associated *Vibrio* strains (17,18). Furthermore, the here reported cell specificity of AIP56 is relevant when considering the potential biotechnological/biomedical use of these toxins, either to inactivate NF-κB in situations where this transcription factor plays an important role or to deliver therapeutic proteins or antigens into the cytosol of myeloid cells.

## 4 Materials and Methods

### 4.1 Production and fluorescence labeling of recombinant proteins

Full-length His-tagged AIP56 (AIP56) and his-tagged AIP56 metalloprotease mutant (AIP56^AAIVAA^), corresponding to a full-length inactive version of the toxin were expressed and purified as previously described (6). Briefly, the sequence encoding AIP56 from the MT1415 strain was cloned in pET28a(+) in frame with a C-terminal His-tag and expressed in *E. coli* BL21 (DE3) cells at 17 °C. The mutant was generated by site directed mutagenesis using QuickChange Site-Directed Mutagenesis Kit (Stratagene) following manufacturer’s instructions. Induced bacterial cells were lysed by sonication, centrifuged, and the recombinant AIP56 was purified from the supernatant using nickel-affinity chromatography (Ni-NTA agarose, ABT) followed by size exclusion chromatography (Superose 12 10/300 GL, GE Healthcare). Inactive AIP56 was purified from inclusion bodies by metal affinity chromatography under denaturing conditions, refolded and followed by size exclusion chromatography. Purified proteins batches were analyzed by SDS-PAGE and purities, determined by densitometry of Coomassie blue-stained gels, were ≥ 90%. Recombinant AIP56 or inactive AIP56 labelled with Alexa Fluor 488 were prepared using the Molecular Probes® Alexa Fluor protein labelling kit (Thermo Fisher Scientific), following the manufacturer’s instructions, but with higher toxin:Alexa488 ratio (57 nM:1 vial), as the recommended ratio inhibited AIP56-488 cell binding and NF-κB p65 cleavage.

### 4.2 Determination of recombinant protein concentration

Protein concentrations were determined by spectrophotometry by measuring absorbance at 280 nm using NanoDrop 1000 (Thermo Fisher Scientific), considering the extinction coefficients and the molecular weights, calculated by the ProtParam tool (http://www.expasy.org/tools/protparam.htmL).

### 4.3 Fish

Sea bass (*Dicentrarchus labrax*) were purchased from commercial hatcheries, and maintained in 600 L seawater aquaria. Water temperature was maintained at 20 ± 1 °C, salinity at 23-28‰ and the photoperiod was 14 h light:10 h dark. Water quality was maintained with mechanical and biological filtration and ozone-disinfection and the fish were fed with commercial pellets (Skretting), adjusting the food intake to fish size and water temperature, according to the supplier’s recommendations.

### 4.4 Mice

C57BL/6 mice were bred and housed at the IBMC/i3S animal facility. They were fed sterilized food and water ad libitum. Four-week-old male mice were used exclusively to derive macrophages from bone marrow. To collect spleens, femurs and tibias, mice were euthanized by CO_2_ followed by cervical dislocation. To collect blood, mice were anesthetized with isoflurane and, after confirmation of the inexistence of reflexes, cardiac puncture was performed. After terminal blood collection, cervical dislocation was done.

### 4.5 Fish neutrophils

A suspension of UV-killed *Phdp* MT1415 virulent strain (OD 600 nm of 0.9), obtained as previously reported (9), was injected ip in sea bass (100 μl per fish) in order to induce inflammation and enrich the peritoneal leukocytes (PL) in neutrophils. After 6 h, fish were injected with 500 µL sea bass PBS (sbPBS; 50 mM phosphate buffer pH 7.2 + 184 mM NaCl) containing 10 µg/mL AIP56-488 or the same volume of sbPBS. After 15 min, PL were collected following a procedure described elsewhere (22). Cells were centrifuged (5 min, 250 × *g*, 4 ºC) and distributed in a 96-well plate (U-bottom). Dead cells were labeled with FVD (eFLuorTM 506 fix viability dye 1:1000 in sbPBS) for 20 min on ice in the dark, centrifuged as above and fixed with 1% paraformaldehyde (PFA) at room temperature (RT) for 15 min. Cells were washed and filtered into FACS tubes. Live neutrophils were selected by differential gating (see section 4.14. bellow).

### 4.6 Fish lymphocytes

Lymphoid cells were obtained from thymus or spleen of sea bass (88.4 - 107.3 g body weight). Fish were euthanized by immersion in 0.06% (v/v) 2-fenoxyethanol, followed by exsanguination, and the organs were removed and placed in isolation medium (L-15 medium (Gibco) adjusted to 322 mOsm and supplemented with 2% fetal bovine serum (FBS, Gibco), 1% penicillin/streptomycin (P/S, Gibco) and 20 U/mL heparin (Braun)). Cell suspensions were obtained by macerating the organs through a 100 µm nylon mesh with isolation medium. Cell suspension was carefully added onto Percoll gradient (45% and 31% Percoll) and centrifuged (45 min, 400 × *g*, 4 °C, without break). The leukocytes remaining at the interface were collected and washed twice with culture medium (L-15 medium adjusted to 322 mOsm and supplemented with 10% fetal bovine serum, 1% P/S and 20 U/mL heparin). The cells were then adjusted to a density of 3 × 10^6^ cells/mL in culture medium and plated in 96-well plate with U-bottom (100 µL/well). Lymphocytes were selected through flow cytometry by differential gating (see section 4.14. bellow).

### 4.7 Isolation of splenocytes, bone marrow, peritoneal and blood cells from mice

Splenocytes were obtained by macerating spleens through a 100 µm nylon mesh with complete DMEM (cDMEM: supplemented with 10% FBS, 10 mM L-glutamine, 10 mM HEPES, 1 mM sodium pyruvate and 1% P/S (all from Gibco)). Then, in the case of splenocytes that were used for cell sorting, red blood cells (RBC) were eliminated by incubation of the cell pellet with 5 mL of RBC lysis buffer (BioLegend) for 5 min at RT. Cell suspension was washed with PBS and centrifuged (5 min, 280 g, 4ºC).

To collect the bone marrow, muscle from mice femurs and tibias was removed, the extremities of the bones cut and the bone marrow cavities flushed with 5 mL of ice-cold cDMEM per bone using a syringe with a 26G needle.

Recruitment of neutrophils into the peritoneal cavity was achieved by the ip injection of 2.5 mL 3% thioglycolate (50). After 6 h, mice were euthanized as described in section 4.4. and 5 mL RPMI with 5% FBS were injected in the peritoneal cavity and left for 1 min before collecting the cells by aspiration with a Pasteur pipette.

Blood was collected by cardiac puncture with a 25G needle on a 1 mL syringe coated with heparin and placed in an Eppendorf with 20 µL heparin. RBC were eliminated by incubation of blood with 10 mL of RBC lysis buffer for 5 min at RT. Cell suspension was washed with 20 mL of PBS and centrifuged (250 × *g*, 10 min, 4 °C).

Splenocytes, bone marrow, peritoneal and blood cells were centrifuged (280 × *g*, 10 min, 4 °C), resuspended in cDMEM and plated at a density of 1 × 10^6^ cells/well in U-bottom 96-well plates for internalization assays. Instead, the pellet of splenocytes to be sorted was resuspended in 3 mL of cDMEM.

### 4.8 Differentiation and polarization of mouse bone marrow derived macrophages

Mouse macrophages were derived from the bone marrow, following an established protocol (51). Bone marrow cells were obtained as previously described, but using Hanks’ balanced salt solution (HBSS) (Gibco) to flush the bone marrow cavities, instead of cDMEM. Cell suspension was centrifuged (250 × *g*, 10 min, 4 °C) and the resulting pellet was resuspended in cDMEM with 10% L929 cell conditioned medium (LCCM) as a source of M-CSF (52). Cells were incubated overnight on a cell culture dish at 37 °C in a 7% CO_2_ atmosphere to remove fibroblasts. Non-adherent cells were collected in HBSS, centrifuged, resuspended in cDMEM with 10% LCCM and plated at a density of 5 × 10^5^ cells/well in a 24-well plate with 1 mL per well. After 3 days, 10% (v/v) LCCM was added to each well and, at day 7, the medium was renewed. At day 10, the medium was removed and cDMEM with cytokines was added to the wells to polarize the primary macrophages. Five µg/mL of LPS and 300 U/mL of IFNγ were used to induce M1-like phenotype, 20 ng/mL of IL-4 was added to promote M2-like differentiation, and cDMEM alone was used to achieve M0. Mouse M0, M1-like and M2-like macrophages were used after 48 h incubation at 37 °C in a 7% CO_2_ atmosphere.

### 4.9 Isolation of human monocytes and lymphocytes

Peripheral blood mononuclear cells (PBMCs) were isolated from human buffy coats from healthy blood donors. Buffy coats were diluted 1:1 in PBS and gently layered over Histopaque^®^-1077 (Sigma). After centrifuging (30 min, 400 × *g*, RT, without break), the whitish ring containing PBMCs was collected and washed with PBS (10 min, 250 g, RT). Cells were resuspended in RBC lysis buffer, incubated for 10 min at RT to eliminate red blood cells and washed with PBS. Magnetic Activated Cell Sorting (MACS) system was used to isolate the desired leukocytes, using nano-sized magnetic beads. Basically, cell pellet was resuspended in PBS and washed in MACS buffer (PBS with 0.5% BSA and 2 mM EDTA). MACS buffer (530 µL) and CD14 beads (70 µL) (MicroBeads human 2 mL 130-050-201) were added per each 100 × 10^6^ PBMCs and incubated for 20 min on ice. PBMCs were washed twice in MACS buffer and resuspended in the same buffer at a density of 100 × 10^6^ cells/mL.

To isolate monocytes, the suspension of PBMCs with CD14 beads was gravity run through Low-Shear (LS) columns pre-equilibrated with MACS buffer. The run-through was collected to isolate lymphocytes, as described below. The CD14 cells remaining in the column were then flushed, washed with PBS, and (i) either resuspended in RPMI 1640 (Gibco) supplemented with 10% FBS, 1% non-essential amino acids, 1 mM sodium pyruvate and 1% kanamycin (all from Gibco) and plated at a density of 0.4 × 10^6^ cells/well in 96-well plate for flow cytometry or 1 × 10^6^ cells/well in 24-well plate for p65 cleavage assays, or (ii) used for obtaining macrophages and DCs as described in sections 4.10 and 4.11, respectively.

Lymphocytes were isolated from the run through of the suspension of PBMCs with CD14 beads, containing CD14-negative cells. Per each 100 × 10^6^ cells from this suspension, 560 µL MACS buffer and 140 µL CD19 beads (MicroBeads human 2mL 130-050-301) were added following by incubation for 20 min on ice. The suspension was then run through LS columns, the flow through collected for acquiring a population enriched in T-cells and the B-cells flushed from the column. After washing with PBS, B- and T-lymphocytes were resuspended in RPMI 1640 supplemented as referred above and plated as described for monocytes.

### 4.10 Polarization of human macrophages

Macrophages were generated following a procedure described elsewhere (53). Monocytes isolated as referred above, were resuspended in RPMI 1640 supplemented with 10% FBS, 1% P/S and 50 ng/mL of M-CSF (ImmunoTools, Friesoythe, Germany) and plated at a density of 0.3 × 10^6^ cells/well in 24-well plates with glass coverslips. Cells were incubated for 7 days at 37 °C with 5% CO_2_. Medium was replaced by fresh RPMI 1640 supplemented with 10% FBS and 1% P/S. After 3 days, the medium was replaced by fresh RPMI 1640 supplemented with 10% FBS, 1% P/S to maintain M0 phenotype or with the same medium supplemented with 100 ng/mL of LPS (Sigma-Aldrich, St. Louis, MO, USA) and 25 ng/mL of IFNγ (ImmunoTools, Friesoythe, Germany) to polarize into M1-like macrophages or with 20 ng/mL of IL-4 to polarize into M2-like macrophages. Cells were incubated for 24-48 h at 37 °C, 5% CO_2_ before use.

### 4.11 Generation of human dendritic cells

Dendritic cells were generated as described elsewhere (54). Isolated monocytes were resuspended in RPMI 1640 supplemented with 10% FBS, 1% non-essential amino acids, 1 mM sodium pyruvate, 1% kanamycin, 50 ng/mL IL-4 and 50 ng/mL GM-CSF, plated at a density of 3 × 10^6^ cells/well in 6-well plates and incubated for 4 days at 37 °C with 5% CO_2_. Half of the medium was removed and replaced by fresh RPMI 1640 supplemented as above, but containing 150 ng/mL IL-4 and 150 ng/mL GM-CSF. After 7 days, the medium was discarded and cells were harvested with PBS by pipetting the cultured cells out of the 6-well plate into a tube. Cells were centrifuged (10 min, 250 × *g*, RT), resuspended in RPMI supplemented with 10% FBS, 1% non-essential amino acids, 1 mM sodium pyruvate and 1% kanamycin and plated at a density of 0.4 × 10^6^ cells/well in 96-well plate (U-bottom) for flow cytometry or 0.5–1 × 10^6^ cells/well in 24-well plate for p65 cleavage assays.

### 4.12 Human neutrophils

Human leukocytes were isolated from whole peripheral blood of healthy volunteers collected into EDTA-collection tubes and slowly agitated. Each subject signed a written informed consent before blood collection. To eliminate erythrocytes, 10 mL of pre-warmed (37 °C) RBC lysis buffer were added per each 500 µL of blood, followed by immediate vortexing and incubation for 10 min at RT. After washing (350 × *g*, 5 min, RT) twice with PBS, the cell pellet was resuspended in the same volume of RPMI 1640 supplemented with 10% FBS as the volume of blood used initially, and 100 µL-aliquots were distributed in wells of a 96-well plate (U-bottom). Neutrophils were identified through flow cytometry by differential gating (see section 4.14. bellow).

### 4.13 AIP56-488 internalization assays

To assess AIP56 internalization, sea bass lymphocytes, mouse and human leukocytes were incubated for 15 min on ice with 89 nM AIP56-488 in the respective medium (see above). For sorting of AIP56-488+ B-cells, the same procedure was followed in mouse splenocytes, but using the fluorescent-labelled inactive AIP56. Cells were centrifuged (3 min, 250 × *g*, 4 °C), the supernatant discarded and the pellet resuspended in the same medium without toxin and incubated for 15 min at 22 °C (sea bass lymphocytes) or 30 min at 37 °C (human and mouse cells) to allow toxin internalization. Cells were washed twice with sbPBS (sea bass lymphocytes) or commercial PBS (Gibco) (mouse and human cells). Mouse cells were incubated with FVD (1:1000 in commercial PBS) for 20 min on ice in the dark to label dead cells, centrifuged (3 min, 250 g, 4 °C) and the supernatant discarded. Sea bass lymphocytes were labeled with the anti-IgM monoclonal antibody WDI3 (34) (1:50 in sbPBS, 1 h on ice) followed by a Cyanine 5 (Cy5) fluorescent goat anti-mouse IgG secondary antibody (1:200 in sbPBS, 20 min on ice in the dark), both in sea bass FACS buffer (sbPBS + 2% FBS). Human and mouse cells were labelled with fluorescent antibodies (Table 1) diluted in FACS buffer (PBS + 2% FBS) for 20 min on ice in the dark in a final volume of 50 μL except for splenocytes to be sorted as AIP56-488+ B-cells, which were incubated in a final volume of 2 mL because the cell density was higher. Cells were washed with its respective FACS buffer, fixed with 1% PFA for 15 min at RT and washed again with FACS buffer. In the case of mBMDM, cells were incubated for 10 min at RT with permeabilization buffer (0.1% saponin in FACS buffer), centrifuged and incubated for 30 min at 4 °C with intracellular antibody anti CD206-APC in permeabilization buffer. Cells were resuspended in its corresponding FACS buffer and filtered through a 35 µm nylon mesh cell strainer into flow cytometry tubes. Propidium iodide (PI; 5 µg/mL; PercP-Cy5.5) was added exclusively to sea bass lymphocytes to access cell viability (instead of FVD). Fluorescence was analyzed by flow cytometry. Both AIP56-488^+^ and inactive AIP56-488^-^ B-cells from mouse splenocytes were isolated into 15 mL tubes through the BD FACSARIA™ II cell sorter, centrifuged (10 min, 250 × *g*, 4ºC), resuspended in 400 µL of cDMEM without FBS and distributed into 2 wells of 96-well plates (U-bottom) at a density of ≈ 1 × 10^6^ cells/well to perform NF-κB p65 cleavage assay.

**Table 1.**
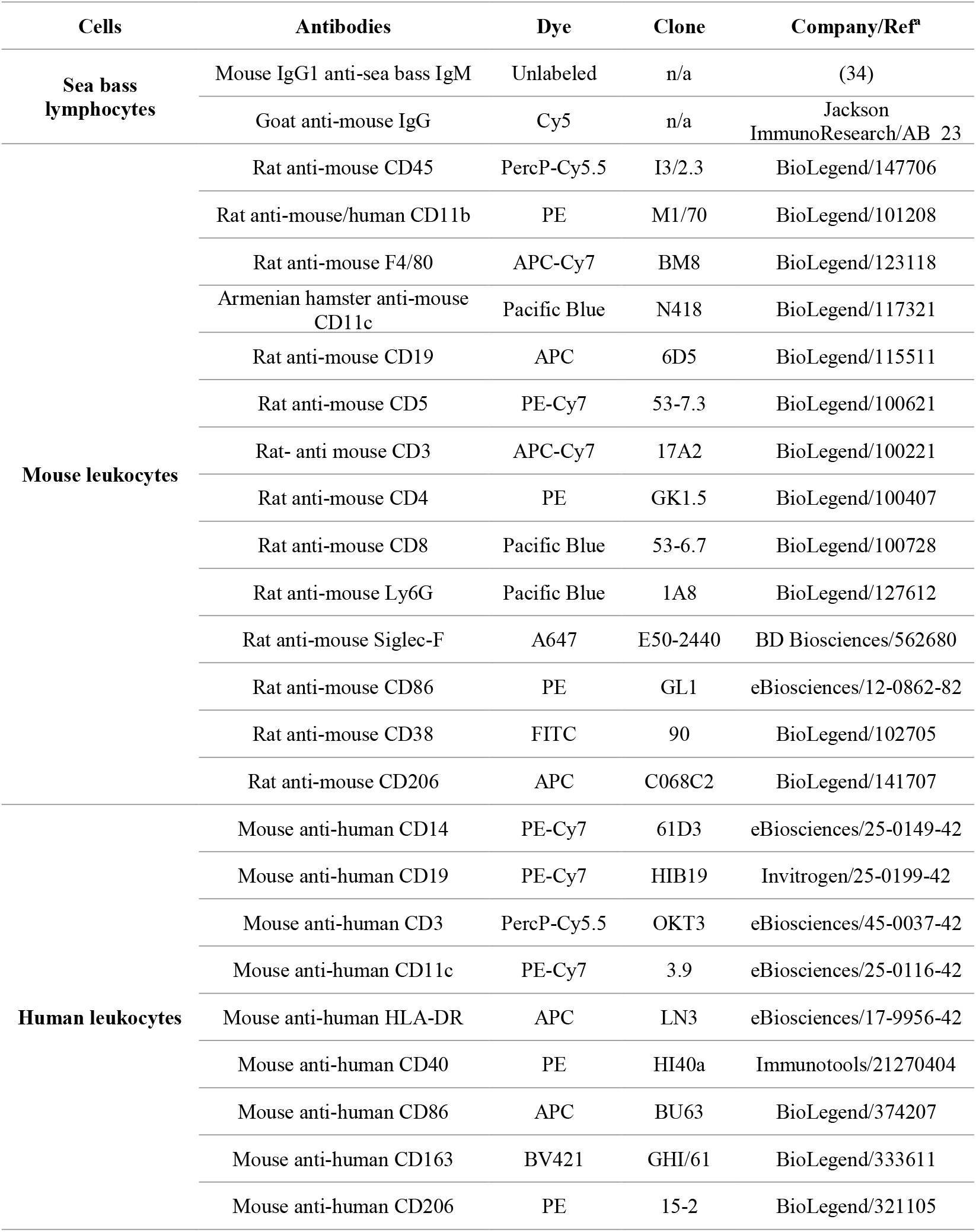
Antibodies used for flow cytometry.

### 4.14 Flow cytometry

Flow cytometry data were acquired on a BD FACSCanto™ II (BD Biosciences, San Jose, CA). Ten thousand events per sample for sea bass and human cells and twenty thousand events per sample for mouse cells were analyzed at medium flow rate. All data were analyzed using FlowJo™ software v10.6.1 (San Diego, CA).

#### Sea bass

Sea bass peritoneal granulocytes and lymphoid cells from spleen and thymus (Figure S1) were first selected by differential gating based on their side scatter area (SSC-A; cell complexity) and forward scatter area (FSC-A; cell size) profile, followed by forward scatter high (FSC-H) and FSC-A profile for elimination of duplets. As neutrophils are the major cell population in sea bass peritoneal cells 6 h after injection of UV-killed MT1415 (22,55), the granulocytes selected based on high SSC-A and FSC-A were assumed as neutrophils and macrophages were identified as intermediate SSC-A and high FSC-A. Then, the viable neutrophils or macrophages were identified as FVD^-^ cells. To confirm the identity of sea bass neutrophils and macrophages, the selected populations were sorted separately (BD FACSARIA™ II cell sorter) into 1.5 mL tubes at a density of approximately 2 × 10^5^ cells/tube. Sorted cells were used to perform cytospins, which were then stained for peroxidase using the Antonow technique (which specifically labels neutrophils (22,56), followed by heamacolor (Figure S1). Regarding lymphoid cells, viable B-cells were selected from splenic lymphocytes by the specific IgM marker WDI3 (WDI3^+^ cells) (25,34) combined with a Cy5 fluorescent goat anti-mouse IgG secondary antibody and PI (PI^−^ cells) gating. WDI3^−^ and PI^−^ thymic lymphoid cells were considered viable putative thymocytes, since around 74.9% of the thymic IgM^−^ cells are thymocytes (24–26).

#### Mouse

Mouse leukocytes were first selected through SSC-A vs. FSC-A and FSC-H vs FSC-A, followed by selection of viable mature leukocytes for the specific leukocyte marker CD45 (CD45^+^ positive cells) and viable dye FVD^−^ negative cells. Then, in mouse bone marrow samples (Figure S2), neutrophils were identified as CD11b^+^ and Ly6G^+^ cells and eosinophils as CD11b^+^ and Siglec-F^+^ cells. Within CD11b^+^ and Ly6G^−^ cells, macrophages were identified as F4/80^+^ and CD11b^high^, and monocytes as F4/80^+^ and CD11b^intermediate^. In mouse spleen samples (Figure S2), B-cells were identified as CD19^+^ lymphocytes with B1-cells identified as CD5^+^, whereas B2-cells were assumed as CD5^−^. The CD19^−^ population was used to select SSC-A^−^ and F4/80^+^ cells for identification of spleen macrophages (F4/80^+^; CD11b^high^) and monocytes (F4/80^+^; CD11b^intermediate^). CD11b^+^/F4/80^+^/CD11c^+^ cells were categorized as DCs. After selecting the viable spleen leukocytes, T-cells (CD3^+^) were also identified and then, helper T-cells (CD4^+^ and CD8^−^) and cytotoxic T-cells (CD8^+^ and CD4^−^) were selected. In mouse peritoneal and blood samples (Figure S2), neutrophils were identified as CD11b^+^ and Ly6G^+^ cells and macrophages were also identified in peritoneal cells as F4/80^+^ and CD11b^high^.

The mBMDM M0, M1-like and M2-like phenotypes were confirmed by analyzing the expression of specific cell markers through flow cytometry (Figure S3). mBMDM were first selected through SSC-A vs FSC-A and singlets were identified through FSC-H vs FSC-A plot. Then, CD86, CD38 and CD206 were analyzed. Mouse M1-like macrophages were confirmed by a higher expression of CD86 and CD38 than M0 and M2-like, whereas M2-like phenotype was confirmed by a higher expression of CD206 than in M0 and M1-like cells.

BD FACSARIA™ II cell sorter (BD Biosciences, San Jose, CA) was used to identify and isolate mouse AIP56-488^+^ and AIP56^-^488^−^ B-cells (Figure S4). First, mouse leukocytes were selected by differential gating based on SSC-A vs FSC-A and duplets were eliminated though FSC-H vs FSC-A plot. B-cells were identified as CD19^+^. Then, AIP56-488^+^ and AIP56-488^−^ B-cells were isolated. Around 2 million events of AIP56-488^+^ B-cells and 4 million events of AIP56-488^−^ B-cells were sorted.

#### Human

Regarding human monocyte derived macrophages (Figure S5), after assessing the viable subset of cells based on their SSC-A vs FSC-A profile and eliminating the duplets (FSC-H vs FSC-A), macrophages were identified as CD40^+^ and CD86^+^, with higher expression of these markers in M1-like followed by M2-like and M0. CD163 was very low in M0 and M2-like and absent in M1-like. Besides that, M2-like phenotype was confirmed through the higher expression of CD206 than M0 and M1-like, because of M2-like polarization with IL-4. The phenotype of the other human myeloid and lymphoid cells used was confirmed as follows (Figure S6): neutrophils from peripheral blood were selected based on the SSC-A^high^ vs FSC-A^high^ profile after eliminating duplets (FSC-H vs FSC-A), considering that 95% of the total pool of circulating granulocyte are neutrophils (57,58); monocytes were CD14^+^; moDCs were HLA-DR^+^ and CD11c^+^; B-cells were identified as CD19^+^ and T-cells as CD3^+^.

### 4.15 NF-κB cleavage assay

Human cells (macrophages, monocytes, moDCs and lymphocytes), mBMDM (M0, M1-like and M2-like) and mouse AIP56-488^+^ B-cells or AIP56-488^-^ B-cells were left untreated or incubated for 15 min on ice with 179 nM of AIP56, followed by a 4 h (human monocytes, moDCs and lymphocytes and mouse cells) or 6 h (macrophages) incubation at 37 °C in 5% CO_2_. Cells were washed and lysed in SDS-PAGE loading buffer (50 mM Tris-HCl pH 8.8, 2% SDS, 0.05% bromophenol blue, 10% glycerol, 2 mM EDTA and 100 mM DTT). Samples were heated at 95 °C for 5 min, subjected to 10% SDS-PAGE using Laemmli discontinuous buffer system (59) and transferred into nitrocellulose membranes (GE Healthcare Life science, Buckinghamshire, UK). Staining with Ponceau S was used to verify loading and transfer efficiency. Membranes were then blocked with 5 % (w/v) skimmed powder milk in TBS-T (tris-buffered saline, 0.1 % Tween) for 1 h at RT followed by incubation with an anti-NF-κB p65 rabbit antibody (sc-372, Santa Cruz Biotechnology) diluted 1:3000 in blocking solution. After 3 washes of 5 min with TBS-T, membranes were incubated for 1 h with alkaline phosphatase conjugated secondary anti-rabbit antibody (A9919, Sigma Aldrich) (human cells and mBMDM) or HRP conjugated secondary anti-rabbit antibody (AP311, the Binding Site) (mouse AIP56-488^+^ B-cells), both secondary antibodies were diluted 1:10000 in blocking solution. After 3 washes, immunoreactive bands were detected using nitroblue tetrazolium–5-bromo-4-chloro-3-indolylphosphate (NBT/BCIP) (Promega, Madison, WI, USA), for human cells and mBMDM, or with ECL Dura (34075, Thermo Scientific™) for mouse AIP56-488^+^ B-cells.

### 4.16 Statistical analysis

Data were analyzed through one-way analysis of variance. Since the results were expressed as percentages, the arcsine transformation was used. As homogeneity of variances was not observed, the Kruskal-Wallis non-parametric test was used.. Although there were statistically significant results across groups (conditions), when the pairwise Bonferroni correction was applied for multiple comparisons, it did not yield significant results in a vast majority of the comparisons. This can be explained by the low number of individuals per condition (n=3). Since the figures by themselves are quite explanatory, we decided not to present the statistical results bearing in mind the limitation just mentioned.

## 7 Institutional Review Board Statement

This work was carried out in accordance with European and Portuguese legislation for the use of animals for scientific purposes (Directive 2010/63/EU; Decreto-Lei 113/2013). The study using sea bass was approved by the ORBEA (Animal Welfare and Ethics Body) of i3S and was licensed by Direcção-Geral de Alimentação e Veterinária (DGAV), the Portuguese authority for animal protection (Lic. 0421/000/000/2021). Human buffy coats from healthy blood donors were obtained from Centro Hospitalar São João (CHSJ) upon request which was approved by CHSJ Ethics Committee for Health (References 259 and 260/11) in compliance with the Declaration of Helsinki ethical principles.

## 8 Informed Consent Statement

All the subjects signed an informed consent before blood donation. Buffy coats were provided anonymized.

## 9 Acknowledgments

The authors acknowledge the support of the i3S Scientific Platform TraCy - Translational Cytometry Unit and the i3S Animal facility.

## 10 Data Availability Statement

The original contributions presented in the study are included in the article and supplementary materials. Further inquiries can be directed to the corresponding authors.

## References

1. Romalde JL. Photobacterium damselae subsp. piscicida: an integrated view of a bacterial fish pathogen. International Microbiology 2002 5:1 (2002) 5:3–9. doi: 10.1007/S10123-002-0051-6

2. Zorrilla I, Balebona MC, Moriñigo MA, Sarasquete C, Borrego JJ. Isolation and characterization of the causative agent of pasteurellosis, Photobacterium damsela ssp. piscicida, from sole, Solea senegalensis (Kaup). J Fish Dis (1999) 22:167–172. doi: 10.1046/J.1365-2761.1999.00157.X

3. Liu PC, Lin JIY, Lee KK. Virulence of Photobacterium damselae subsp. piscicida in cultured cobia Rachycentron canadum. J Basic Microbiol (2003) 43:499–507. doi: 10.1002/JOBM.200310301

4. do Vale A, Silva MT, dos Santos NMS, Nascimento DS, Reis-Rodrigues P, Costa-Ramos C, Ellis AE, Azevedo JE. AIP56, a novel plasmid-encoded virulence factor of Photobacterium damselae subsp. piscicida with apoptogenic activity against sea bass macrophages and neutrophils. Mol Microbiol (2005) 58:1025–1038. doi: 10.1111/j.1365-2958.2005.04893.x

5. do Vale A, Pereira C, R Osorio C, M S dos Santos N. The Apoptogenic Toxin AIP56 Is Secreted by the Type II Secretion System of Photobacterium damselae subsp. piscicida. Toxins (Basel) (2017) 9:368. doi: 10.3390/toxins9110368

6. Silva DS, Pereira LMG, Moreira AR, Ferreira-Da-Silva F, Brito RM. The Apoptogenic Toxin AIP56 Is a Metalloprotease A-B Toxin that Cleaves NF-kb P65. PLoS Pathog (2013) 9:1003128. doi: 10.1371/journal.ppat.1003128

7. Silva MT, do Vale A, dos Santos NMS. Secondary necrosis in multicellular animals: An outcome of apoptosis with pathogenic implications. Apoptosis (2008) 13:463–482. doi: 10.1007/S10495-008-0187-8

8. do Vale A, Costa-Ramos C, Silva A, Silva DSP, Gärtner F, dos Santos NMS, Silva MT. Systemic macrophage and neutrophil destruction by secondary necrosis induced by a bacterial exotoxin in a Gram-negative septicaemia. Cell Microbiol (2007) 9:988–1003. doi: 10.1111/J.1462-5822.2006.00846.X

9. do Vale A, Marques F, Silva MT. Apoptosis of sea bass (Dicentrarchus labrax L.) neutrophils and macrophages induced by experimental infection with Photobacterium damselae subsp. piscicida. Fish Shellfish Immunol (2003) 15:129–144. doi: 10.1016/S1050-4648(02)00144-4

10. Pereira LMG, Pinto RD, Silva DS, Moreira AR, Beitzinger C, Oliveira P, Sampaio P, Benz R, Azevedo JE, dos Santos NMS, et al. Intracellular trafficking of AIP56, an NF-κB-cleaving toxin from Photobacterium damselae subsp. piscicida. Infect Immun (2014) 82:5270–5285. doi: 10.1128/IAI.02623-14

11. Lisboa J, Pereira C, Pinto RD, Rodrigues IS, Pereira LMG, Pinheiro B, Oliveira P, Pereira PJB, Azevedo JE, Durand D, et al. Unconventional structure and mechanisms for membrane interaction and translocation of the NF-κB-targeting toxin AIP56. Nature Communications 2023 14:1 (2023) 14:1–16. doi: 10.1038/s41467-023-43054-z

12. Turco MM, Sousa MC. The Structure and Specificity of the Type III Secretion System Effector NleC Suggest a DNA Mimicry Mechanism of Substrate Recognition. Biochemestry (2014) doi: 10.1021/bi500593e

13. Baruch K, Gur-Arie L, Nadler C, Koby S, Yerushalmi G, Ben-Neriah Y, Yogev O, Shaulian E, Guttman C, Zarivach R, et al. Metalloprotease type III effectors that specifically cleave JNK and NF-κB. EMBO Journal (2011) 30:221–231. doi: 10.1038/EMBOJ.2010.297

14. Yen H, Ooka T, Iguchi A, Hayashi T, Sugimoto N, Tobe T. NleC, a Type III Secretion Protease, Compromises NF-κB Activation by Targeting p65/RelA. PLoS Pathog (2010) 6:e1001231. doi: 10.1371/JOURNAL.PPAT.1001231

15. Gilmore TD, Wolenski FS. NF-κB: Where did it come from and why? Immunol Rev (2012) 246:14–35. doi: 10.1111/j.1600-065X.2012.01096.x

16. Rodrigues IS, Pereira LMG, Lisboa J, Pereira C, Oliveira P, dos Santos NMS, do Vale A. Involvement of Hsp90 and cyclophilins in intoxication by AIP56, a metalloprotease toxin from Photobacterium damselae subsp. piscicida. Sci Rep (2019) 9:1–13. doi: 10.1038/s41598-019-45240-w

17. Liang K, Islam MT, Hussain N, Winkjer NS, Im MS, Rowe LA, Tarr CL, Boucher Y. Draft Genome Sequences of Eight Vibrio sp. Clinical Isolates from across the United States That Form a Basal Sister Clade to Vibrio cholerae. Microbiol Resour Announc (2019) 8: doi: 10.1128/MRA.01473-18

18. Islam MT, Liang K, Orata FD, Im MS, Alam M, Lee CC, Boucher YF. Vibrio tarriae sp. nov., a novel member of the Cholerae clade. Int J Syst Evol Microbiol (2022) 72: doi: 10.1099/IJSEM.0.005571

19. Verster KI, Cinege G, Lipinszki Z, Magyar LB, Kurucz É, Tarnopol RL, Ábrahám E, Darula Z, Karageorgi M, Tamsil JA, et al. Evolution of insect innate immunity through domestication of bacterial toxins. Proc Natl Acad Sci U S A (2023) 120:e2218334120. doi: 10.1073/PNAS.2218334120/SUPPL_FILE/PNAS.2218334120.SM01.AVI

20. Piot N, Gisou van der Goot F, Sergeeva OA. Harnessing the Membrane Translocation Properties of AB Toxins for Therapeutic Applications. Toxins 2021, Vol 13, Page 36 (2021) 13:36. doi: 10.3390/TOXINS13010036

21. Freitas IL, Teixeira A, Loureiro I, Lisboa J, Saraiva A, dos Santos NMS, do Vale A. Susceptibility of Sea Bream (Sparus aurata) to AIP56, an AB-Type Toxin Secreted by Photobacterium damselae subsp. piscicida. Toxins (Basel) (2022) 14: doi: 10.3390/TOXINS14020119

22. do Vale A, Afonso A, Silva MT. The professional phagocytes of sea bass (Dicentrarchus labrax L.): cytochemical characterisation of neutrophils and macrophages in the normal and inflamed peritoneal cavity. Fish Shellfish Immunol (2002) 13:183–198. doi: 10.1006/FSIM.2001.0394

23. Esteban MÁ, Muñoz J, Meseguer J. Blood Cells of Sea Bass (Dicentrarchus labrax L.). Flow Cytometric and Microscopic Studies. Anat Rec (2000) 258:80–89. doi: 10.1002/(SICI)1097-0185(20000101)258:1

24. Scapigliati G, Mazzini M, Mastrolia L, Romano N, Abelli L. Production and characterisation of a monoclonal antibody against the thymocytes of the sea bass Dicentrarchus labrax(L.) (Teleostea, Percicthydae). Fish Shellfish Immunol (1995) 5:393–405. doi: 10.1006/FSIM.1995.0039

25. dos Santos NMS, Romano N, de Sousa M, Ellis AE, Rombout JHWM. Ontogeny of B and T cells in sea bass (Dicentrarchus labrax, L.). Fish Shellfish Immunol (2000) 10:583–596. doi: 10.1006/FSIM.2000.0273

26. Picchietti S, Buonocore F, Guerra L, Belardinelli MC, De Wolf T, Couto A, Fausto AM, Saraceni PR, Miccoli A, Scapigliati G. Molecular and cellular characterization of European sea bass CD3ε + T lymphocytes and their modulation by microalgal feed supplementation. Cell Tissue Res 1:3. doi: 10.1007/s00441-020-03347-x

27. Freitas IL, Teixeira A, Loureiro I, Lisboa J, Saraiva A, dos Santos NMS, do Vale A. Susceptibility of Sea Bream (Sparus aurata) to AIP56, an AB-Type Toxin Secreted by Photobacterium damselae subsp. piscicida. Toxins (Basel) (2022) 14: doi: 10.3390/TOXINS14020119

28. Ma RY, Black A, Qian BZ. Macrophage diversity in cancer revisited in the era of single-cell omics. Trends Immunol (2022) 43:546–563. doi: 10.1016/J.IT.2022.04.008

29. Lendeckel U, Venz S, Wolke C. Macrophages: shapes and functions. ChemTexts (2022) 8:12. doi: 10.1007/S40828-022-00163-4

30. Martinez FO, Gordon S. The M1 and M2 paradigm of macrophage activation: Time for reassessment. F1000Prime Rep (2014) 6:1–13. doi: 10.12703/P6-13

31. Franken L, Schiwon M, Kurts C. Macrophages: Sentinels and regulators of the immune system. Cell Microbiol (2016) 18:475–487. doi: 10.1111/cmi.12580

32. Singhal A, Kumar S. Neutrophil and remnant clearance in immunity and inflammation. Immunology (2022) 165:22–43. doi: 10.1111/IMM.13423

33. Nauseef WM. Human neutrophils ≠ murine neutrophils: Does it matter? Immunol Rev (2022) doi: 10.1111/IMR.13154

34. dos Santos NMS, Taverne N, Taverne-Thiele AJ, de Sousa M, Rombout JHWM. Characterisation of monoclonal antibodies specific for sea bass (Dicentrarchus labrax L.) IgM indicates the existence of B cell subpopulations. Fish Shellfish Immunol (1997) 7:175–191. doi: 10.1006/FSIM.1996.0073

35. Stegelmeier AA, van Vloten JP, Mould RC, Klafuric EM, Minott JA, Wootton SK, Bridle BW, Karimi K. Myeloid Cells during Viral Infections and Inflammation. Viruses (2019) 11: doi: 10.3390/V11020168

36. McDaniel MM, Meibers HE, Pasare C. Innate control of Adaptive Immunity and Adaptive Instruction of Innate Immunity: Bi-Directional flow of information. Curr Opin Immunol (2021) 73:25. doi: 10.1016/J.COI.2021.07.013

37. Pidwill GR, Gibson JF, Cole J, Renshaw SA, Foster SJ. The Role of Macrophages in Staphylococcus aureus Infection. Front Immunol (2020) 11: doi: 10.3389/FIMMU.2020.620339

38. Wang C, Yu X, Cao Q, Wang Y, Zheng G, Tan TK, Zhao H, Zhao Y, Wang Y, Harris DCH. Characterization of murine macrophages from bone marrow, spleen and peritoneum. BMC Immunol (2013) 14: doi: 10.1186/1471-2172-14-6

39. Louwe PA, Badiola Gomez L, Webster H, Perona-Wright G, Bain CC, Forbes SJ, Jenkins SJ. Recruited macrophages that colonize the post-inflammatory peritoneal niche convert into functionally divergent resident cells. Nature Communications 2021 12:1 (2021) 12:1–15. doi: 10.1038/s41467-021-21778-0

40. Wang J, Kubes P. A Reservoir of Mature Cavity Macrophages that Can Rapidly Invade Visceral Organs to Affect Tissue Repair. Cell (2016) 165:668–678. doi: 10.1016/J.CELL.2016.03.009

41. Bou Ghosn EE, Cassado AA, Govoni GR, Fukuhara T, Yang Y, Monack DM, Bortoluci KR, Almeida SR, Herzenberg LA, Herzenberg LA. Two physically, functionally, and developmentally distinct peritoneal macrophage subsets. Proc Natl Acad Sci U S A (2010) 107:2568–2573. doi: 10.1073/PNAS.0915000107/SUPPL_FILE/PNAS.200915000SI.PDF

42. Bassler K, Schulte-Schrepping J, Warnat-Herresthal S, Aschenbrenner AC, Schultze JL. The Myeloid Cell Compartment-Cell by Cell. Annu Rev Immunol (2019) 37:269–293. doi: 10.1146/ANNUREV-IMMUNOL-042718-041728/CITE/REFWORKS

43. Geert Wiegertjes F, Philip Elks M. Fish Macrophages. Principles of Fish Immunology: From Cells and Molecules to Host Protection (2022) 203–227. doi: 10.1007/978-3-030-85420-1_6

44. Tsai JM, Shoham M, Fernhoff NB, George BM, Marjon KD, McCracken MN, Kao KS, Sinha R, Volkmer AK, Miyanishi M, et al. Neutrophil and monocyte kinetics play critical roles in mouse peritoneal adhesion formation. Blood Adv (2019) 3:2713. doi: 10.1182/BLOODADVANCES.2018024026

45. Popi AF, Longo-Maugéri IM, Mariano M. An overview of B-1 cells as antigen-presenting cells. Front Immunol (2016) 7:138. doi: 10.3389/FIMMU.2016.00138/BIBTEX

46. Popi AF. B-1 phagocytes: the myeloid face of B-1 cells. Ann N Y Acad Sci (2015) 1362:86–97. doi: 10.1111/NYAS.12814

47. Tangye SG. To B1 or not to B1: that really is still the question! Blood (2013) 121:5109–5110. doi: 10.1182/BLOOD-2013-05-500074

48. Kuby. Immunology. 7th ed. W. H. Freeman and Company. (2012). 81–87 p. doi: 10.1007/s13398-014-0173-7.2

49. Smith FL, Baumgarth N. B-1 cell responses to infections. Curr Opin Immunol (2019) 57:23–31. doi: 10.1016/J.COI.2018.12.001

50. Oliveira L, Silva MC, Gomes AP, Santos RF, Cardoso MS, Nóvoa A, Luche H, Cavadas B, Amorim I, Gärtner F, et al. CD5L as a promising biological therapeutic for treating sepsis. doi: 10.1038/s41467-024-48360-8

51. Gomes MS, Fernandes SS, Cordeiro J V, Gomes SS, Vieira A, Appelberg R. Engagement of Toll-like receptor 2 in mouse macrophages infected with Mycobacterium avium induces non-oxidative and TNF-independent anti-mycobacterial activity. Eur J Immunol (2008) 38:2180–2189. doi: 10.1002/eji.200737954

52. Englen MD, Valdez YE, Lehnert NM, Lehnert BE. Granulocyte/macrophage colony-stimulating factor is expressed and secreted in cultures of murine L929 cells. J Immunol Methods (1995) 184:281–283. doi: 10.1016/0022-1759(95)00136-X

53. Loureiro JP, Cruz MS, Cardoso AP, Oliveira MJ, Macedo MF. Human iNKT Cells Modulate Macrophage Survival and Phenotype. Biomedicines (2022) 10:1723. doi: 10.3390/BIOMEDICINES10071723/S1

54. Pereira CS, Pérez-Cabezas B, Ribeiro H, Maia ML, Cardoso MT, Dias AF, Azevedo O, Ferreira MF, Garcia P, Rodrigues E, et al. Lipid antigen presentation by CD1b and CD1d in lysosomal storage disease patients. Front Immunol (2019) 10:435588. doi: 10.3389/FIMMU.2019.01264/BIBTEX

55. Ferreira IA, Peixoto D, Losada AP, Quiroga MI, Vale A do, Costas B. Early innate immune responses in European sea bass (Dicentrarchus labrax L.) following Tenacibaculum maritimum infection. Front Immunol (2023) 14: doi: 10.3389/FIMMU.2023.1254677/FULL

56. Afonso A, Lousada S, Silva J, Ellis AE, Silva MT. Neutrophil and macrophage responses to inflammation in the peritoneal cavity of rainbow trout Oncorhynchus mykiss. A light and electron microscopic cytochemical study. Dis Aquat Organ (1998) 34:27–37.

57. Krouse JH. Introduction to Allergy. Managing the Allergic Patient (2008) 1–17. doi: 10.1016/B978-141603677-7.50005-4

58. Prinyakupt J, Pluempitiwiriyawej C. Segmentation of white blood cells and comparison of cell morphology by linear and naïve Bayes classifiers. Biomed Eng Online (2015) 14:1–19. doi: 10.1186/S12938-015-0037-1/TABLES/8

59. Laemmli UK. Cleavage of Structural Proteins during the Assembly of the Head of Bacteriophage T4. Nature 1970 227:5259 (1970) 227:680–685. doi: 10.1038/227680a0

