## Supplementary figures for "AIP56, an AB toxin secreted by *Photobacterium damselae* subsp. *piscicida*, has tropism for myeloid cells"

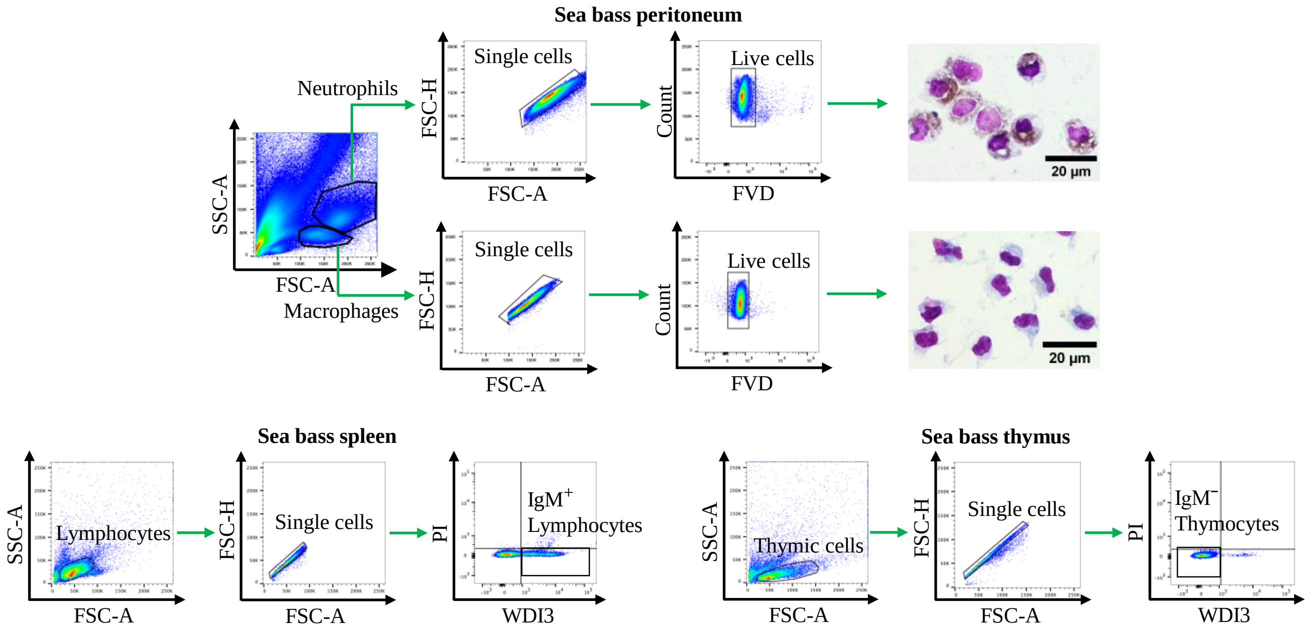


**Figure S1.** Gating strategy applied to identify sea bass neutrophils and macrophages from peritoneum and $IgM^+$ lymphocytes (B-cells) and IgM- cells (putative thymocytes) from sea bass spleen and thymus samples, respectively, after Percoll gradient. Neutrophils were assessed based on their morphology (FSC-A vs SSC-A) considering that the analyzed samples were from inflamed peritoneal cavities after 6 h injection of UV-killed MT1415, where neutrophils are the majority of the granulocytes. Macrophages from the same cavities were also assessed based on FSC-A vs SSC-A and analyzed as control. After selecting viable neutrophils and macrophages, cells were sorted and cytospins were performed to confirm the identity of the selected cells. Hemacolor technique was performed and Antonow was used for peroxidase detection to stain neutrophils (brown), allowing to distinguish between these two cells types. Lymphoid cells were selected based on their morphology (SSC-A vs FSC-A) and IgM expression. Fixable viability dye (FVD) and propidium iodide (PI) were used to identify viable neutrophils and lymphoid cells, respectively.

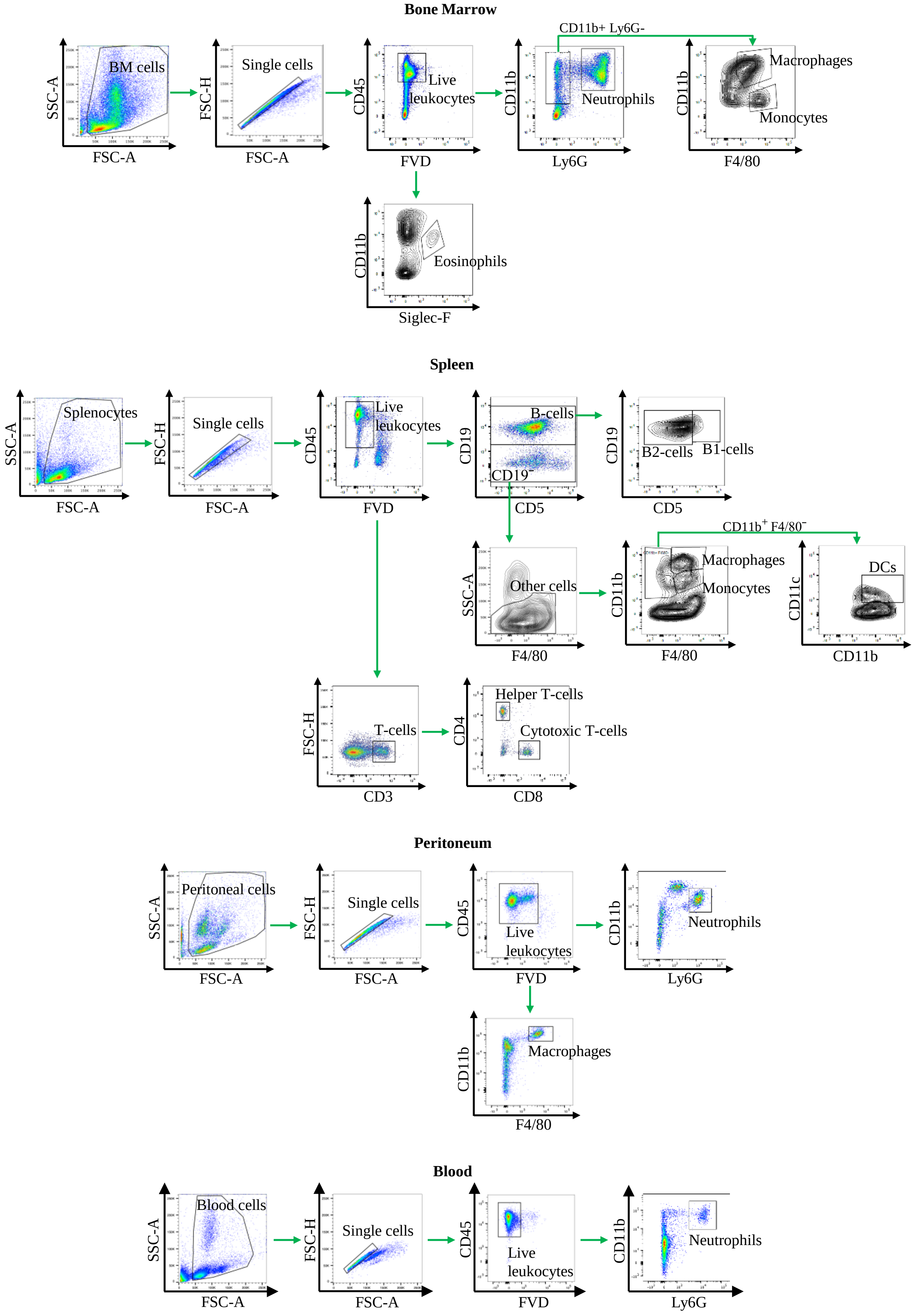


**Figure S2.** Gating strategy applied to identify different mouse leukocytes from bone marrow (BM), spleen, peritoneum (inflamed with thioglycolate for 6h) and blood samples, after identifying general cell population by morphological parameters (SSC-A vs FSC-A) and eliminating duplets (FSC-H vs FSC-A).

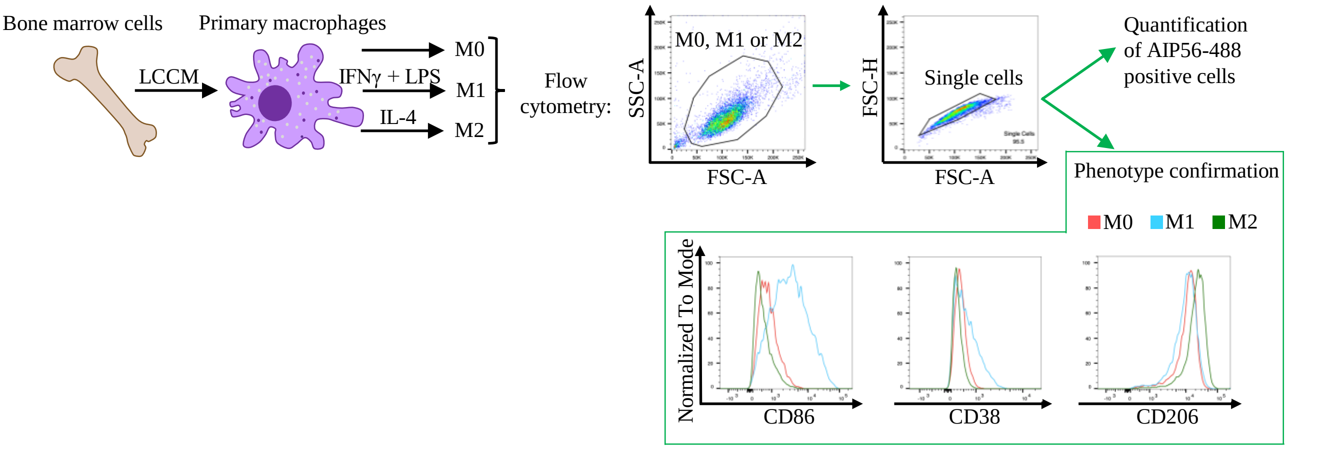


**Figure S3.** Schematic description of mBMDM differentiation and polarization. Representative plots of the gating strategy applied to confirm M0, M1-like and M2-like phenotype, after identifying general cell population by morphological parameters (SSC-A vs FSC-A) and eliminating duplets (FSC-H vs FSC-A), based on their expression of CD86, CD38 and CD206, respectively.

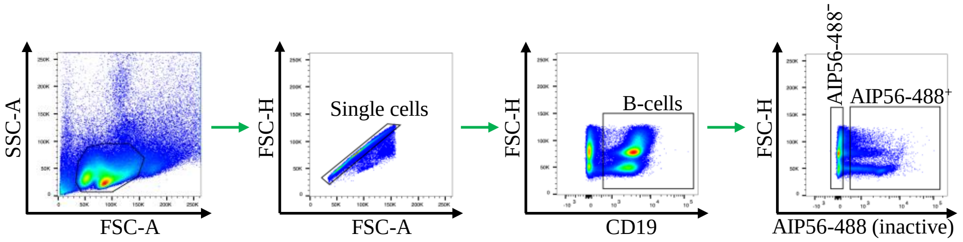


**Figure S4.** Gating strategy applied to isolate mouse AIP56-488$^{+}$ and AIP56-488$^{-}$ B-cells from spleen, after identifying general cell population by morphological parameters (SSC-A vs FSC-A), eliminating duplets (FSC-H vs FSC-A) and selecting B-cells (CD19$^{+}$).

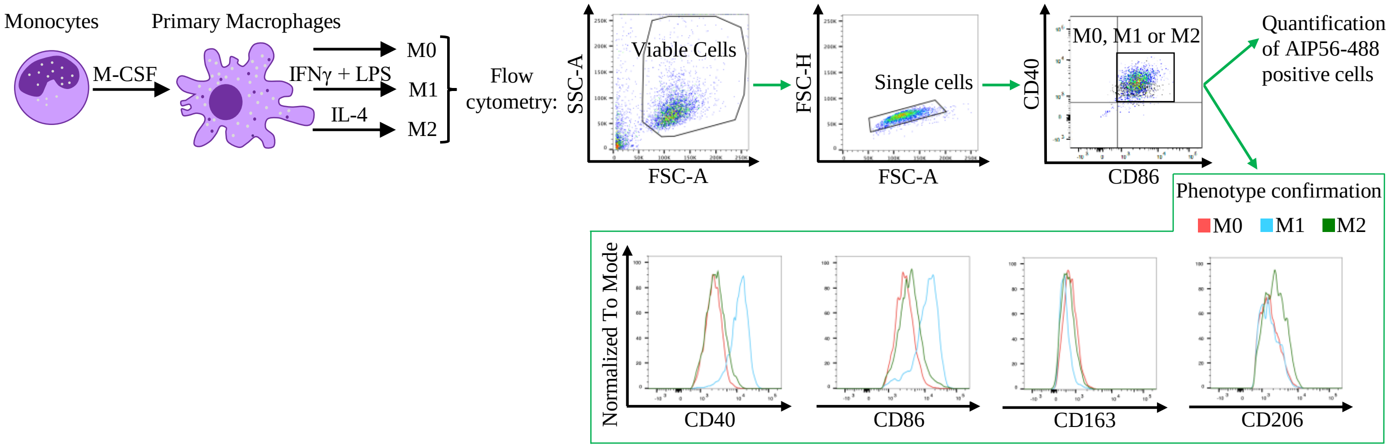


**Figure S5.** Schematic description of human monocytes differentiation into macrophages and their polarization. Representative plots of gating strategy applied to identify human macrophages, after selecting the general cell population through morphological parameters (SSC-A vs FSC-A) and eliminating duplets (FSC-H vs FSC-A). Macrophages subtypes (M0, M1-like and M2-like) were confirmed based on their expression of CD40, CD86, CD163 and CD206, respectively.

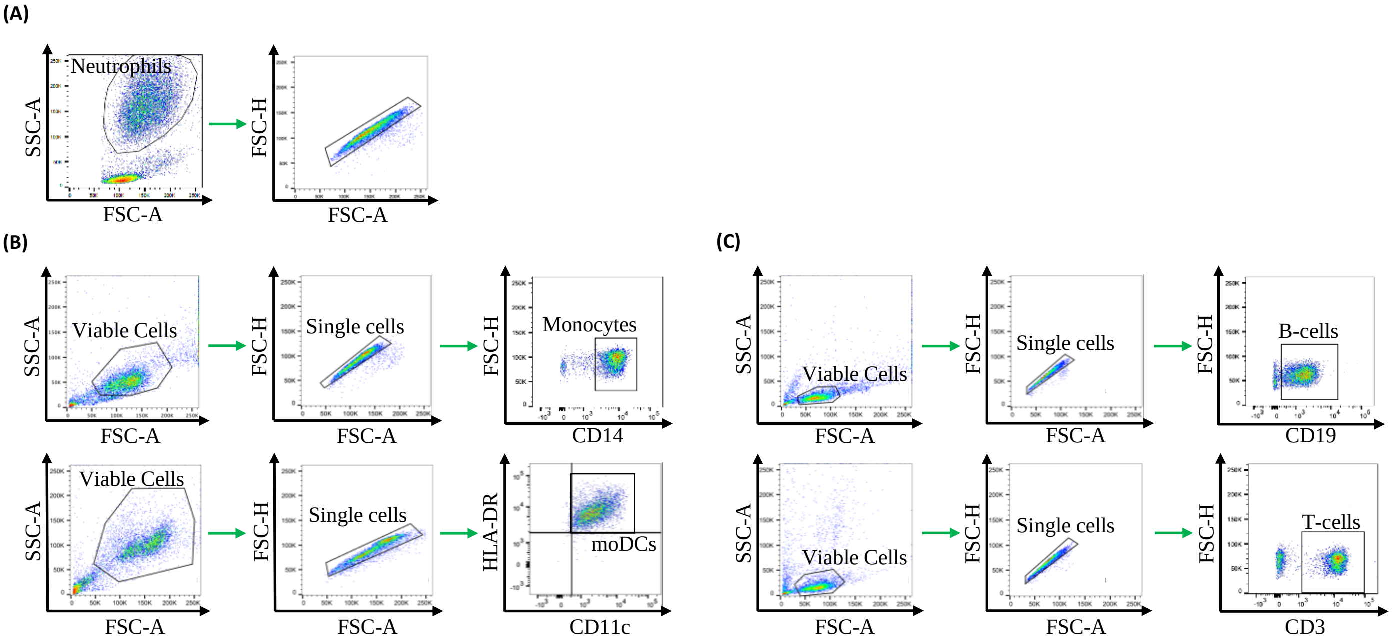


**Figure S6.** Gating strategy applied to identify different and viable human leukocytes. **(A)** Neutrophils were analyzed from whole peripheral blood, where neutrophils are the majority of the granulocytes. Neutrophils were gated based on morphological parameters (SSC-A vs FSC-A) and then, duplets were eliminated (FSC-H vs FSC-A). **(B)** Monocytes were isolated from buffy coats by MACS system, using CD14 beads and moDCs were derived from these cells. **(C)** The portion of cells that were negatively selected by CD14 beads were divided in B-cells and T-cells also by MACS system, using CD19 beads. **(B, C)** Gating strategy to identify the obtained monocytes, moDCs, B-cells and T-cells after selecting the general cell population through morphological parameters (SSC-A vs FSC-A) and eliminating duplets (FSC-H vs FSC-A).
