## Supplementary tables for "AIP56, an AB toxin secreted by *Photobacterium damselae* subsp. *piscicida*, has tropism for myeloid cells"

**Table S1.** AIP56-488 internalization by sea bass leukocytes.

<table>
<tr><th colspan="3">Sea bass cell populations</th><th colspan="2">% of fluorescent cells</th><th>MFI[1]</th></tr>
<tr><th>Lineage</th><th>Source</th><th>Cell type</th><th>Untreated cells</th><th>AIP56-488 treated cells</th><th>AIP56-488 treated cells</th></tr>
<tr><td rowspan="2">Myeloid</td><td rowspan="2">Peritoneum</td><td>Macrophages</td><td>0.06 ± 0.13</td><td>82.38 ± 9.44</td><td>46.83 ± 16.66</td></tr>
<tr><td>Neutrophils</td><td>0.01 ± 0.02</td><td>13.40 ± 7.25</td><td>2.14 ± 0.90</td></tr>
<tr><td rowspan="2">Lymphoid</td><td>Spleen</td><td>$IgM^+$ Lymphocytes</td><td>0.67 ± 0.87</td><td>2.53 ± 0.43</td><td>1.36 ± 0.09</td></tr>
<tr><td>Thymus</td><td>$IgM^-$ Thymocytes</td><td>0.10 ± 0.07</td><td>0.58 ± 0.44</td><td>1.29 ± 0.19</td></tr>
</table>

[1]MFI (Median Fluorescence Intensity) results are presented as fold-change in fluorescence intensity to the untreated cells (control). Values from each individual experiment are shown in Supplementary Material data sheet 1.

**Table S2.** AIP56-488 internalization by mouse leukocytes.

<table>
<tr><th colspan="4">Mouse cell populations</th><th colspan="2">% of fluorescent cells</th><th>MFI[1]</th></tr>
<tr><th>Lineage</th><th>Source</th><th colspan="2">Cell type</th><th>Untreated cells</th><th>AIP56-488 treated cells</th><th>AIP56-488 treated cells</th></tr>
<tr><td rowspan="13">Myeloid</td><td rowspan="7">BM</td><td colspan="2">Macrophages</td><td>0.31 ± 0.20</td><td>72.83 ± 9.86</td><td>6.47 ± 1.63</td></tr>
<tr><td colspan="2">Monocytes</td><td>0.82 ± 0.46</td><td>25.30 ± 3.29</td><td>2.12 ± 0.14</td></tr>
<tr><td colspan="2">Eosinophils</td><td>0.59 ± 0.56</td><td>3.87 ± 2.44</td><td>1.25 ± 0.12</td></tr>
<tr><td colspan="2">Neutrophils</td><td>0.18 ± 0.04</td><td>3.28 ± 1.17</td><td>1.16 ± 0.37</td></tr>
<tr><td rowspan="3">mBMDM</td><td>M0</td><td>2.93 ± 0.48</td><td>23.50 ± 6.07</td><td>1.67 ± 0.23</td></tr>
<tr><td>M1</td><td>2.08 ± 0.75</td><td>16.70 ± 2.33</td><td>2.07 ± 0.17</td></tr>
<tr><td>M2</td><td>2.13 ± 0.69</td><td>36.60 ± 14.41</td><td>2.13 ± 0.41</td></tr>
<tr><td rowspan="3">Spleen</td><td colspan="2">Macrophages</td><td>0.38 ± 0.65</td><td>85.37 ± 2.57</td><td>8.86 ± 3.12</td></tr>
<tr><td colspan="2">Monocytes</td><td>1.11 ± 1.39</td><td>20.40 ± 4.13</td><td>2.24 ± 0.57</td></tr>
<tr><td colspan="2">DCs</td><td>0.00 ± 0.00</td><td>20.05 ± 6.45</td><td>3.32 ± 0.19</td></tr>
<tr><td rowspan="2">Peritoneum</td><td colspan="2">Macrophages</td><td>1.28 ± 0.19</td><td>98.30 ± 1.31</td><td>28.66 ± 9.41</td></tr>
<tr><td colspan="2">Neutrophils</td><td>0.28 ± 0.13</td><td>8.61 ± 1.72</td><td>0.95 ± 0.28</td></tr>
<tr><td>Blood</td><td colspan="2">Neutrophils</td><td>0.00 ± 0.01</td><td>7.00 ± 5.29</td><td>1.92 ± 0.18</td></tr>
<tr><td rowspan="6">Lymphoid</td><td rowspan="6">Spleen</td><td colspan="2">B-cells</td><td>0.58 ± 0.96</td><td>8.17 ± 0.68</td><td>2.43 ± 0.88</td></tr>
<tr><td colspan="2">B1-cells</td><td>2.83 ± 3.68</td><td>33.85 ± 7.43</td><td>5.76 ± 0.64</td></tr>
<tr><td colspan="2">B2-cells</td><td>0.62 ± 0.74</td><td>10.89 ± 7.23</td><td>2.87 ± 0.41</td></tr>
<tr><td colspan="2">T-cells</td><td>0.09 ± 0.13</td><td>1.99 ± 0.99</td><td>1.19 ± 0.06</td></tr>
<tr><td colspan="2">Tc-cells</td><td>0.00 ± 0.00</td><td>1.02 ± 0.86</td><td>1.16 ± 0.09</td></tr>
<tr><td colspan="2">Th-cells</td><td>0.08 ± 0.08</td><td>1.11 ± 0.01</td><td>1.10 ± 0.02</td></tr>
</table>

[1]MFI (Median Fluorescence Intensity) results are presented as fold-change in fluorescence intensity to the untreated cells (control). Values from each individual experiment are shown in Supplementary Material data sheet 1.

**Table S3.** AIP56-488 internalization by human leukocytes.

| Human cell populations | | % of fluorescent cells | | MFI[1] |
|---|---|---|---|---|
| **Lineage** | **Cell type** | **Untreated cells** | **AIP56-488 treated cells** | **AIP56-488 treated cells** |
| **Myeloid** | M0 | 1.18 ± 0.76 | 91.87 ± 12.02 | 20.98 ± 4.17 |
| | M1 | 0.48 ± 0.32 | 95.19 ± 4.16 | 7.66 ± 4.87 |
| | M2 | 1.62 ± 1.18 | 96.62 ± 3.03 | 25.15 ± 10.87 |
| | Monocytes | 0.07 ± 0.06 | 93.57 ± 8.64 | 9.59 ± 4.44 |
| | moDCs | 0.47 ± 0.25 | 72.20 ± 15.57 | 5.30 ± 2.35 |
| | Neutrophils | 3.76 ± 6.04 | 61.10 ± 27.25 | 2.83 ± 0.91 |
| **Lymphoid** | B-cells | 0.16 ± 0.24 | 7.07 ± 0.85 | 1.50 ± 0.59 |
| | T-cells | 0.05 ± 0.03 | 1.10 ± 0.10 | 1.15 ± 0.10 |

[1]MFI (Median Fluorescence Intensity) results are presented as fold-change in fluorescence intensity to the untreated cells (control). Values from each individual experiment are shown in Supplementary Material data sheet 1.
